# Proteomic insights into the pathophysiology of hypertension-associated albuminuria: Pilot study in a South African cohort

**DOI:** 10.1101/2023.10.30.564666

**Authors:** Melanie A. Govender, Stoyan H. Stoychev, Jean-Tristan Brandenburg, Michèle Ramsay, June Fabian, Ireshyn S. Govender

## Abstract

**Background:** Hypertension is an important public health priority with a high prevalence in Africa. It is also an independent risk factor for kidney outcomes. We aimed to identify potential proteins and pathways involved in hypertension-associated albuminuria by assessing urinary proteomic profiles in black South African participants with combined hypertension and albuminuria compared to those who have neither condition.

**Methods:** The study included 24 South African cases with both hypertension and albuminuria and 49 control participants who had neither condition. Protein was extracted from urine samples and analysed using ultra-high-performance liquid chromatography coupled with mass spectrometry. Data was generated using data-independent acquisition (DIA) and processed using Spectronaut™ 15. Statistical and functional data annotation were performed on Perseus and Cytoscape to identify and annotate differentially abundant proteins. Machine learning was applied to the dataset using the OmicLearn platform.

**Results:** Overall, a mean of 1,225 and 915 proteins were quantified in the control and case groups, respectively. Three hundred and thirty-two differentially abundant proteins were constructed into a network. Pathways associated with these differentially abundant proteins included the immune system (q-value [false discovery rate]=1.4×10^-45^), innate immune system (q=1.1×10^-32^), extracellular matrix (ECM) organisation (q=0.03) and activation of matrix metalloproteinases (q=0.04). Proteins with high disease scores (76–100% confidence) for both hypertension and CKD included angiotensinogen (AGT), albumin (ALB), apolipoprotein L1 (*APOL1*), and uromodulin (UMOD). A machine learning approach was able to identify a set of 20 proteins, differentiating between cases and controls.

**Conclusions:** The urinary proteomic data combined with the machine learning approach was able to classify disease status and identify proteins and pathways associated with hypertension and albuminuria.

## Introduction

Globally, the World Health Organization estimates that more than a billion people have hypertension, a risk factor for adverse cardiovascular and cerebrovascular events and kidney disease (1). Hypertension is a major cause of premature death worldwide, with a disproportionate burden in low- and middle-income countries, largely due to a rising prevalence of risk factors in recent decades (1).

As an upper middle-income country, South Africa has a high prevalence of hypertension, and hypertensive kidney disease is the most common cause of kidney failure documented by the South African Renal Registry (2). Hypertension and kidney disease are inextricably linked, with common pathophysiological denominators that begin with foetal programming in-utero (3, 4). There is strong epidemiological evidence linking low birth weight, as a marker of adverse intrauterine circumstances, to adult hypertension and kidney disease (4, 5). Potential mechanisms include a congenital deficit in nephron number which may arise from changes in DNA methylation, accelerated apoptosis in the developing kidney, changes in renal renin– angiotensin system activity, and an increase in foetal glucocorticoid exposure (4). A decrease in nephron number is associated with compensatory glomerular hypertrophy and an increased susceptibility to the progression of kidney disease (4).

Angiotensin, an endocrine hormone peptide, is a vital part of the renin-angiotensin-aldosterone system, an inter-related endocrine system that plays a significant role in volume and blood pressure control (6). In response to a drop in blood pressure, or sympathetic nerve activity in the kidney, renin is released and cleaves off two amino acids enzymatically to form angiotensin I (ATI), which is cleaved by the angiotensin converting enzyme (ACE) to form angiotensin II (ATII) (6). ATII is the main effector molecule of this system, increasing blood pressure, enhancing renal tubular reabsorption of sodium and water, and stimulating aldosterone release from the adrenal gland (7). In addition to being an effective vasoconstrictor, ATII has also been shown to activate proliferative, pro-inflammatory and pro-fibrotic pathways and stimulates the production of ET-1 resulting in an increase in oxidative stress (7–9). These combined effects of ATII contribute to the development of kidney disease (10–12).

With sustained elevations of blood pressure, the afferent arterioles in the kidney undergo structural changes and hypertrophy, and intraglomerular pressure results in glomerular hypertrophy, hyperfiltration, and albuminuria (13–15). Sustained glomerular hyperfiltration can result in glomerular scarring and irreversible kidney injury (13–17).

Kidney disease is diagnosed using estimated glomerular filtration rate (eGFR) and albuminuria, as each independently confer and increased risk of cardiovascular and all-cause mortality (18). Chronic kidney disease is defined as eGFR <60 mL/min/1.73 m2; and/ or albuminuria, defined as an albumin:creatinine ratio ≥[3.0 mg/mmol (18). Albuminuria may precede the onset of reduced eGFR, which makes it appealing as a potential for early detection and treatment strategies for kidney disease, but this is not a consistent, replicable finding (19, 20). The search for more sensitive markers of early kidney disease has extended to exploration of urinary proteomics with significant progress in the proteomic analysis of biological fluids, in this case, urine.

The past decade has seen omics studies contributing to the diagnosis, therapeutic intervention and prognosis of kidney disease (21–23). Samples derived from urine are non-invasive and easy to collect compared to other body fluids, and can be used to identify genomic, metabolomic, transcriptomic, and proteomic biomarkers that are strongly associated with pathophysiologic mechanisms of disease (24). Proteomic strategies such as mass spectrometry combined with liquid chromatography and capillary electrophoresis are used to identify urine biomarkers involved in early detection of kidney disease and potential disease pathways that provide insight into the pathogenic mechanisms of disease (25–27). One such example is a classifier based on 273 urinary peptides (CKD273) which reliably allows for early detection of kidney disease and is more sensitive than albuminuria in predicting a decline in eGFR (27).

To date, there is one study from South Africa that has investigated the urinary proteome in young adults with hypertension (28). This study found that combining 20 peptides into a single classifier resulted in the separation of normotensive and hypertensive groups with an area under the curve of 0.85 (*P*<0.001) (28). There are no published studies from South Africa investigating the urinary proteome in the setting of hypertension and albuminuria – as an early indicator of kidney injury. In this exploratory study, we hypothesise that urinary proteomic analysis will identify urinary markers of kidney disease in hypertensive individuals with albuminuria and well preserved eGFR.

## Methods

### Study participants

This case-control pilot study is a sub-study of the African Research Kidney Disease (ARK) study, a well characterised population-based cohort study of 2021 adults (20–80 years) of self-identified black ethnicity from Agincourt in the rural Mpumalanga Province, South Africa (29). Demographic, health, and family history information were obtained for all participants, with collection of stored urine and blood samples. From the cohort, ninety participants were selected for proteomic analysis (**Figure 1**).

**Figure 1.**
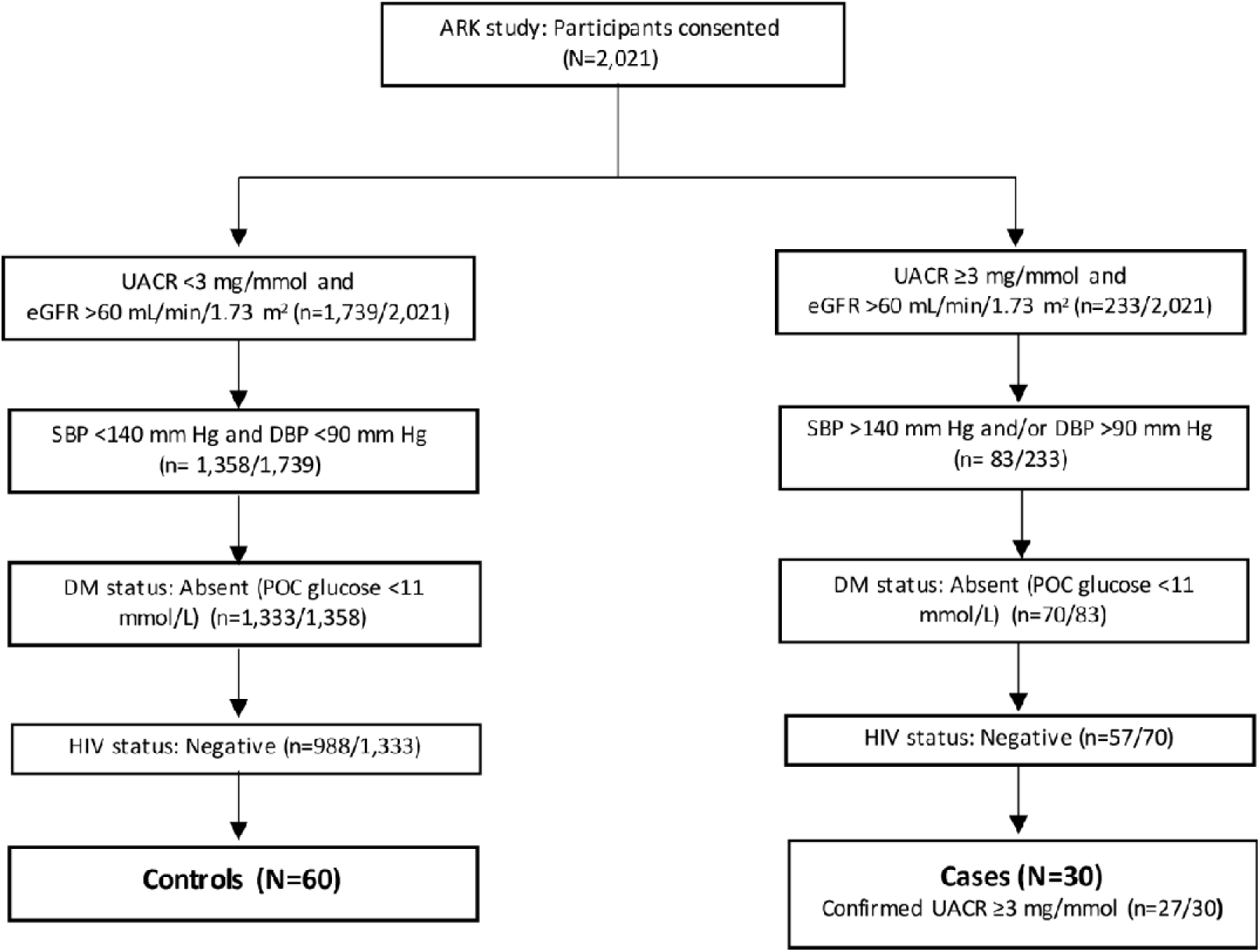
Flow diagram for this case-control study. ARK, African Research Kidney Disease; DBP, diastolic blood pressure; DM, diabetes mellitus; HIV; human immunodeficiency virus; POC, point of care; SBP, systolic blood pressure; UACR, urine albumin-to-creatinine ratio.

Cases were defined as having both albuminuria (urine albumin-to-creatinine ratio [UACR] ≥3 mg/mmol) and hypertension (SBP ≥140 mm Hg and/or diastolic blood pressure [DBP] ≥90 mm Hg, JNC7 criteria). The control group was matched by age and sex. Individuals with other potential causes or consequences of kidney disease, including diabetes mellitus and HIV infection were excluded.

### Urine collection and Measurement of eGFR and UACR

Approximately 20–30 ml urine was collected from ARK study participants. Urine samples were processed and stored at –80°C prior to shipping to the Council for Scientific and Industrial Research laboratory for proteomic testing (30). Serum and urine creatinine was measured using an isotope dilution mass spectrometry traceable Jaffe method (31). The 2009 Chronic Kidney Disease Epidemiology Collaboration (CKD-EPI) creatinine equation without adjustment for African American ethnicity was used to calculate eGFR (ml/min/1.73 m^2^) (32). Immunoturbidimetry was used to measure urine albumin concentration (31). The UACR was calculated (mg/mmol) based on this measurement (31).

### Urinary protein extraction by precipitation

An in-house urinary proteome sample preparation method was used. For each urine sample, protein precipitation was achieved by adding 1,600 µL ice-cold acetone to 400 µL crude urine, stored at -32°C for 1 h. The samples were centrifuged at 12,000 g at 4°C for 1 h. Thereafter, the supernatant was discarded and the pellet was allowed to air dry. The precipitated protein pellet was resuspended in 100 µL 2% sodium dodecyl sulfate solution and sonicated for 5 min (Elmi, Riga, Latvia). Dithiothreitol was added to the solution (final 10 mM), and placed on a 70°C heating block for 15 min, and then on a 40°C heating block for 15 min. The tubes were cooled to room temperature (RT) and iodoacetamide was added for alkylation (final 30 mM) and kept in the dark at RT for 30 min. The protein sample was mixed with an equal volume HILIC binding buffer (30% acetonitrile [MeCN]/200mM Ammonium acetate [NH_4_Ac]) and kept at RT before it was added to a KingFisher™ deep well plate.

### Automated on-bead digestion using MagReSyn^®^ HILIC

Protein samples were digested on-bead using multimode magnetic microparticles (MagReSyn^®^ HILIC, ReSyn Biosciences) in a KingFisher Duo™ system (Thermo Fisher Scientific), as previously described (33, 34), with minor modifications. Briefly, magnetic hydrophilic affinity microparticles (20 μl, 200 μg) were equilibrated in 200 μl 100 mM NH_4_Ac pH 4.5, 15% MeCN. The microparticles were transferred to a well containing the protein-binding buffer solution and mixed for 30 min at RT. The captured proteins were washed twice in 200μl 95% MeCN and transferred to 200μl 25mM ammonium bicarbonate containing 1 μg sequencing grade modified trypsin (Promega, Madison, USA) and mixed for 4 h at 37°C. Finally, beads were washed in 1% trifluoracetic acid to elute any remaining bound peptides. The resulting peptides (pool of digest and eluate) were vacuum dried, resuspended in 2% MeCN, 0.2% FA and quantified using the Pierce™ Peptide Quantification (Thermo Fisher Scientific) assay as per the manufacturer’s instructions.

A project specific system suitability-quality control (PQC) sample was prepared by pooling an equal volume of 16 urinary peptide samples. Additionally, a complex proteome digest was used as a general system suitability assessment. These PQC samples were injected at least once with each batch and analysed and processed together with the study samples.

### Low pH reverse phase liquid chromatography with mass spectrometry (LCMS/MS) data acquisition

Individual participant peptide samples (500 ng, single shot) were analysed using a Dionex UltiMate™ 3000 UHPLC in nanoflow configuration. Samples were inline desalted on an Acclaim PepMap C18 trap column (75 μm × 2 cm; 2 min at 5 μl/min using 2% MeCN/0.2% FA). Trapped peptides were gradient eluted and separated on a nanoEase M/Z Peptide CSH C18 Column (130 Å, 1.7 µm, 75 µm X 250 mm) (Waters) at a flow-rate of 300 nl/min with a gradient of 6–35% over 30 min (A: 0.1% FA; B: 80% MeCN/0.1% FA).

Data was acquired using DIA - or Sequential Window Acquisition of all Theoretical Mass Spectra (SWATH) using a TripleTOF^®^ 6600 mass spectrometer (SCIEX, Massachusetts, USA) (35). Eluted peptides were delivered into the mass spectrometer via a Nanospray^®^ III ion source equipped with a 20 µm Sharp Singularity emitter (Fossil Ion Technology, Madrid, Spain). Source settings were: Curtain gas - 20, Gas 1 - 16, Gas 2 - 0, temperature – 0 (off) and ion spray voltage – 2 600 V.

Data was acquired using 64 MS/MS scans of overlapping sequential precursor isolation windows (variable m/z isolation width, 1 m/z overlap, high sensitivity mode), with a precursor MS scan for each cycle. The accumulation time was 100 ms for the MS1 scan (from 400 to 900 m/z) and 15 ms for each product ion scan (100 to 1 800 m/z) for a 1.06 sec cycle time.

### DIA (SWATH) library generation and data extraction

A spectral library was built (from all patient DIA files) in Spectronaut™ 15 software using default settings with minor adjustments as follows: segmented regression was used to determine the normalised retention time (iRT) in each run; iRTs were calculated as median for all runs; the digestion rule was set as “Trypsin” and modified peptides were allowed; fragment ions between 300 and 1 800 m/z and ions with greater than 3 amino acids were considered; peptides with a minimum 3 and maximum 6 (most intense) fragment ions were accepted. This study specific spectral library was concatenated with an in-house generated urinary proteome spectral library (using Spectronaut™ 15’s “Search Archives” feature).

Raw SWATH (.wiff) data files were converted into Spectronaut™ HTRMS format and analysed using Spectronaut™ 15. The default settings that were used for targeted analysis were: dynamic iRT prediction with correction factor for window 1; mass calibration was set to local; decoy method set as scrambled; the FDR, based on the mProphet approach (36), set at 1% on the precursor peptide and protein group levels; protein inference set to “default” which is based on the ID picker algorithm (37) and global cross-run normalisation on median. The concatenated urinary proteome spectral library (peptides – 20 616, protein groups – 2 604) was used as a reference for targeted data extraction.

Following acquisition, data was curated and filtered for sample preparation and/or LCMS related failures.

### Statistical and functional data annotation

For the table of descriptive data, t-tests and a Chi-squared analysis using STATA were used to compute associations between the case and control group, as appropriate.

Protein abundances were imported into SRplot and log_2_ transformed. Principal component analysis (PCA) based on protein abundances was performed to assess stratification within the cohort. A non-parametric t-test was performed to assess the number of proteins that differed between the case and control group. Statistically significant differentially abundant proteins between the cases and controls were calculated by a two-sided t-test, with a cut-off minimum of 2-fold difference and *P* values adjusted for multiple testing by FDR at 1%.

The differentially abundant proteins were constructed into networks and annotated using Cytoscape and STRING functional enrichment (38). Enrichment was considered statistically significant when corrected for multiple testing by FDR with adjusted q-values <0.05. Additional networks were imported on Cytoscape using public databases for hypertension and CKD with a high confidence score of 0.8 (i.e., 80% confidence) and maximum 200 proteins.

### Machine learning

OmicLearn [v1.4] was used for data analysis, model execution, and creation of plots and charts (39). Machine learning was performed in Python (3.10.12). Spectronaut feature tables were imported via the Pandas package (1.5.3) and manipulated using the Numpy package (1.24.2). The machine learning pipeline was employed using the scikit-learn package (1.2.2). The Plotly (5.9.0) library was used to generate plots and charts. No normalisation on the data was performed. To impute missing values, a Zero-imputation strategy was used. Features were selected using a ExtraTrees (n_trees=100) strategy with a maximum number of 20 features. During training, normalisation and feature selection was individually performed using the data of each split. The XGBoost-Classifier (version: 1.7.4, random_state=23, learning_rate=0.3, min_split_loss=0, max_depth=6, min_child_weight=1) was used for classification.

## Results

Of the initial 90 participants selected for proteomic analyses, 9 were removed due to poor peptide/protein recoveries and 8 were removed due to poor liquid chromatography separation possibly due to incomplete removal of sample specific contamination during sample preparation. Baseline characteristics of the overall ARK study and the 24 case and 49 control participants whose data passed quality control are displayed in **Table 1**.

**Table 1.**
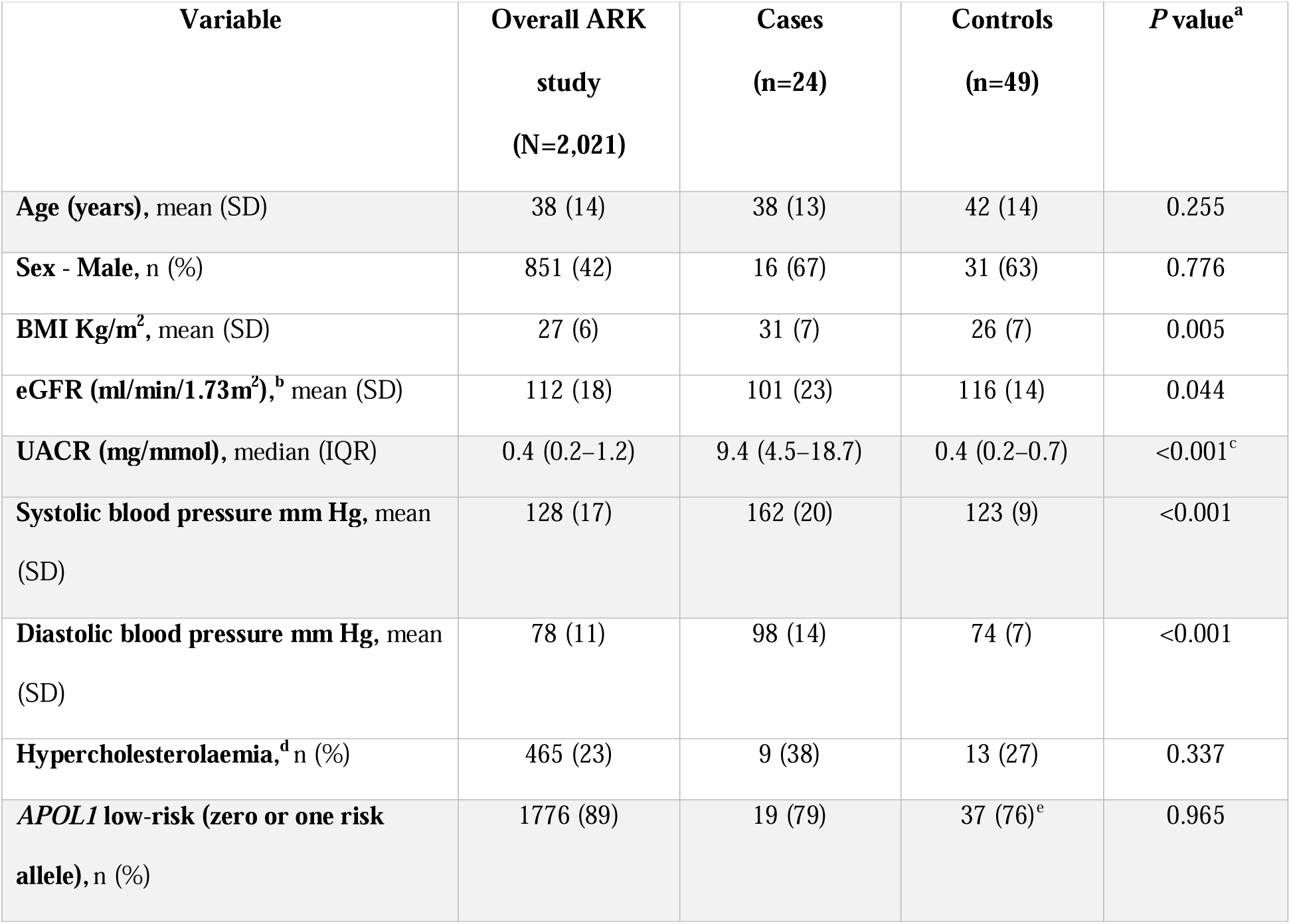
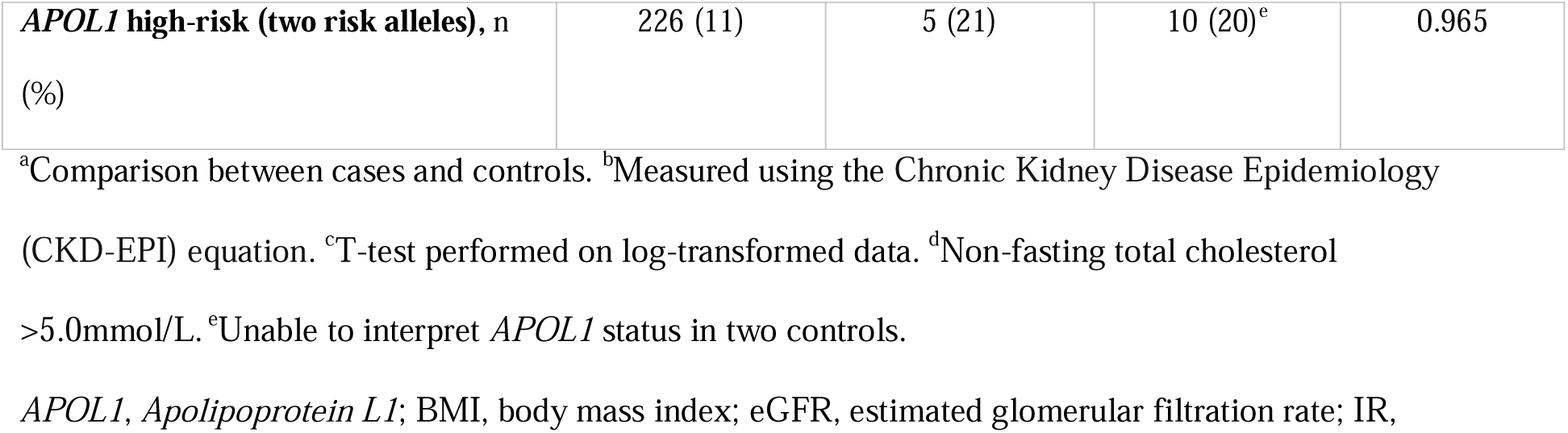
Baseline characteristics of participants in the ARK study and this case-control study.

The age and sex distribution of participants was similar in the case and control groups. The case group had a higher mean body mass index compared to controls, and as expected, blood pressure and UACR differed between the groups. In addition, *apolipoprotein L1* (*APOL1*) allele distribution was similar between cases and controls.

### Project specific system suitability-quality control

The study specific and commercial system suitability quality controls are shown in **(Supplementary Figure 1A–F).** The protein co-efficient of variation (CV) was 11.6% for the Hela digest system suitability assessment and 18.0% for the urine peptide pool over the seven days of analysis (**Supplementary Figure 1A and B)**. Protein, peptide and precursor counts were stable throughout the data acquisition process **(Supplementary Figure 1C and D)**. The correlation plots indicate the system was stable with minimal drift over the course of data acquisition (**Supplementary Figure 1E and F)**.

### Identification of proteins by LCMS analysis

On average, 915 (±264) proteins were quantified in the case group and 1,225 (±227) proteins were quantified in the control group (**Figure 2, Supplementary Figure 2)**. The number of quantified proteins significantly differed between the case and control groups (*P*<0.001).

**Figure 2.**
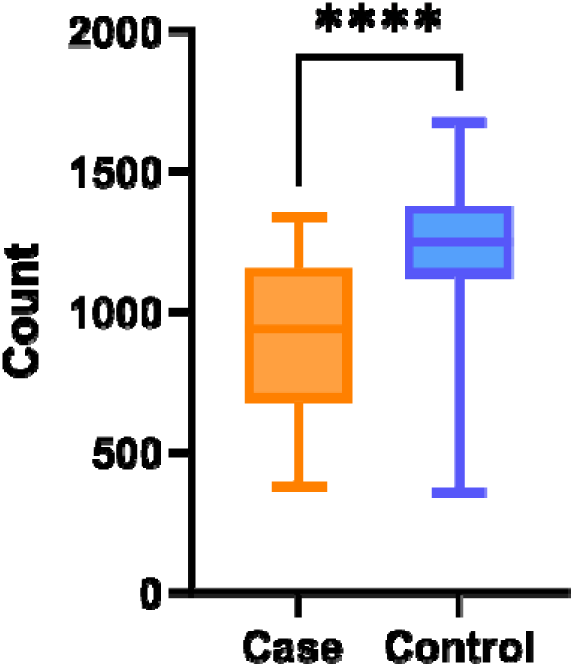
Summary of quantified proteins. The number of proteins quantified post SWATH LCMS analysis for each sample and grouped per condition. LCMS, liquid chromatography with mass spectrometry. ****Denotes *P*<0.001 performed using a non-parametric t-test.

### Principal Component (PC) analysis

PC analysis using protein abundances highlighted differential clustering of the two groups with some overlap (**Figure 3**).

**Figure 3.**
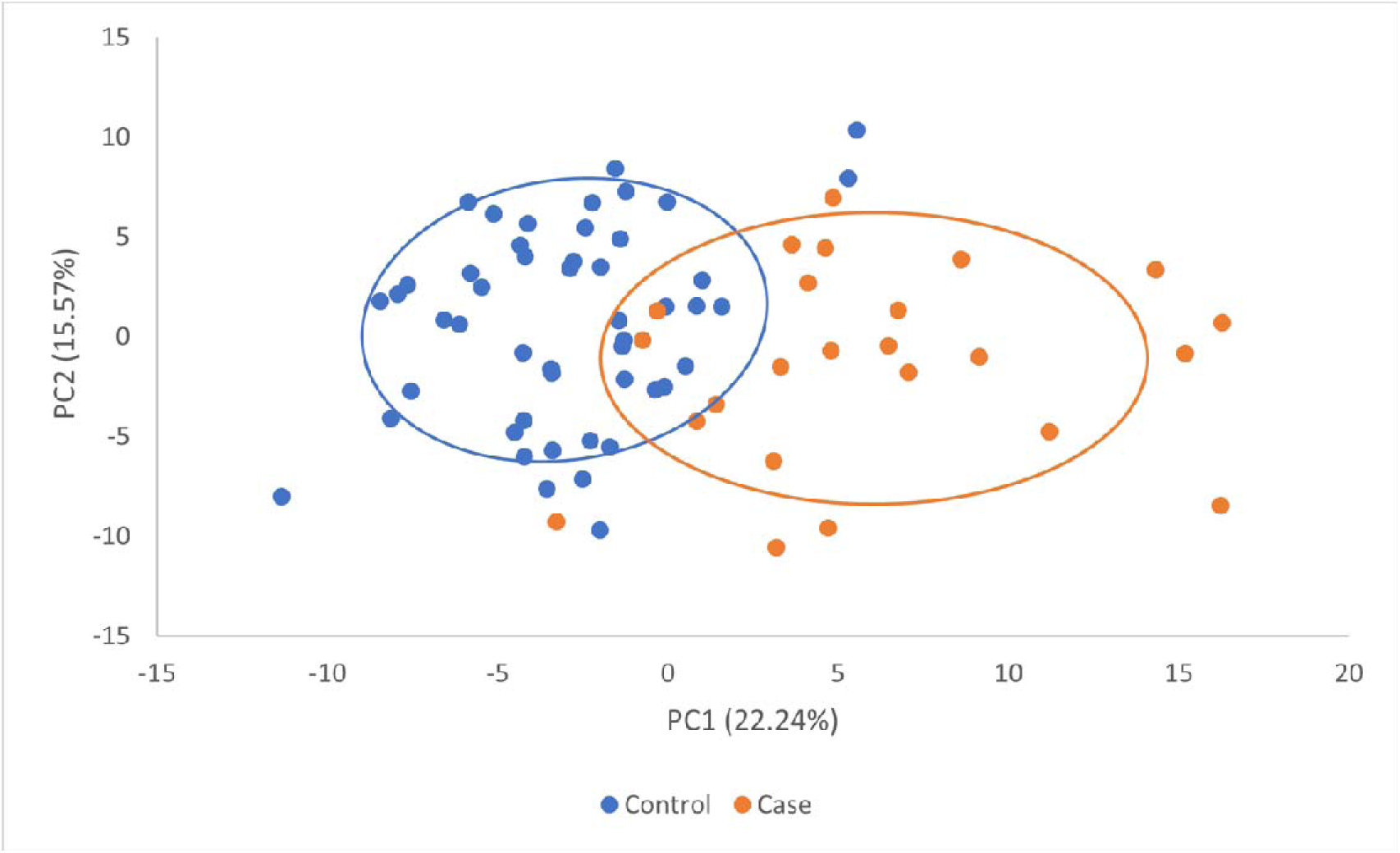
PC (1 and 2) analysis based on protein abundance features. PC analysis using log-transformed protein abundance values indicates clustering of cases and controls. Orange and blue circles represent cases and controls, respectively. PC, principal component.

### Identification of proteins with significantly different abundances between cases and controls

Differentially abundant proteins between the cases and controls are shown in **Figure 4**. Majority of the proteins deemed statistically different in abundances between cases and controls, were found to have lower abundance in the cases. The full list of differential protein abundances is shown in **Supplementary Table 1**.

**Figure 4.**
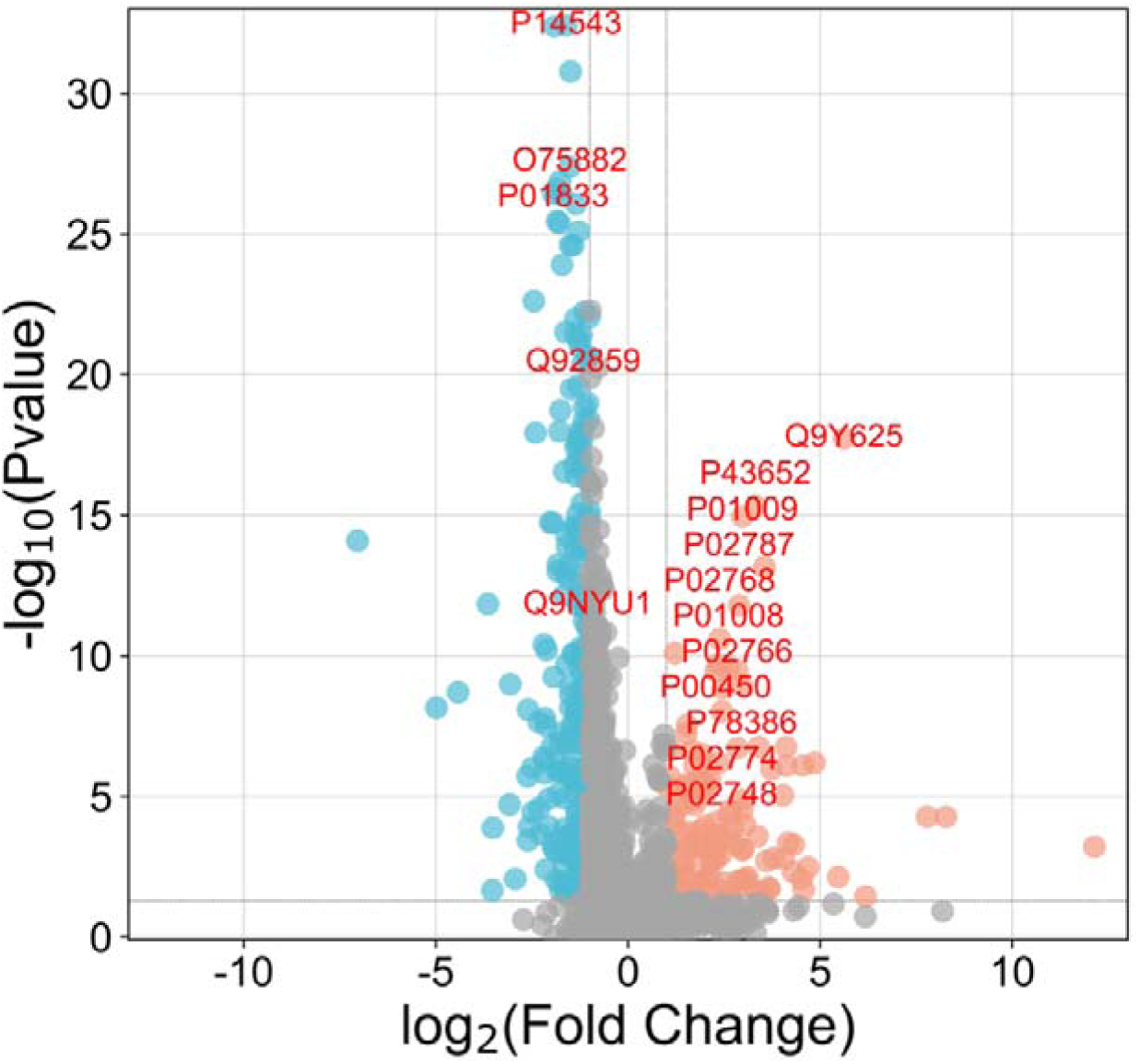
Differential protein abundances between cases and controls illustrated in a volcano plot. Proteins were graphed by fold change (difference) on the x-axis and significance (–log_10_ *P*) on the y-axis using an FDR of 0.01 and a fold-change of 2. Statistical analysis was performed in Perseus. Orange dots indicate proteins with higher abundances in the cases. Blue dots indicate proteins with lower abundances in the cases. The top 20 features of the machine learning algorithm are annotated in red. FDR, false discovery rate.

### Machine learning model

The performance of the machine learning algorithm was evaluated by receiver operating characteristics analysis with an area under the curve of 0.98 and confusion matrix analysis where predicted and actual classification closely matched (**Supplementary Figure 3**). The top 20 features identified from the classifier are shown in **Figures 4 and 5**. When comparing the top 20 features to the list of 332 differentially abundant proteins identified (using a student t-test), 16 were found to overlap and four were unique to the XgBoost algorithm. (**Supplementary Table 2**).

**Figure 5.**
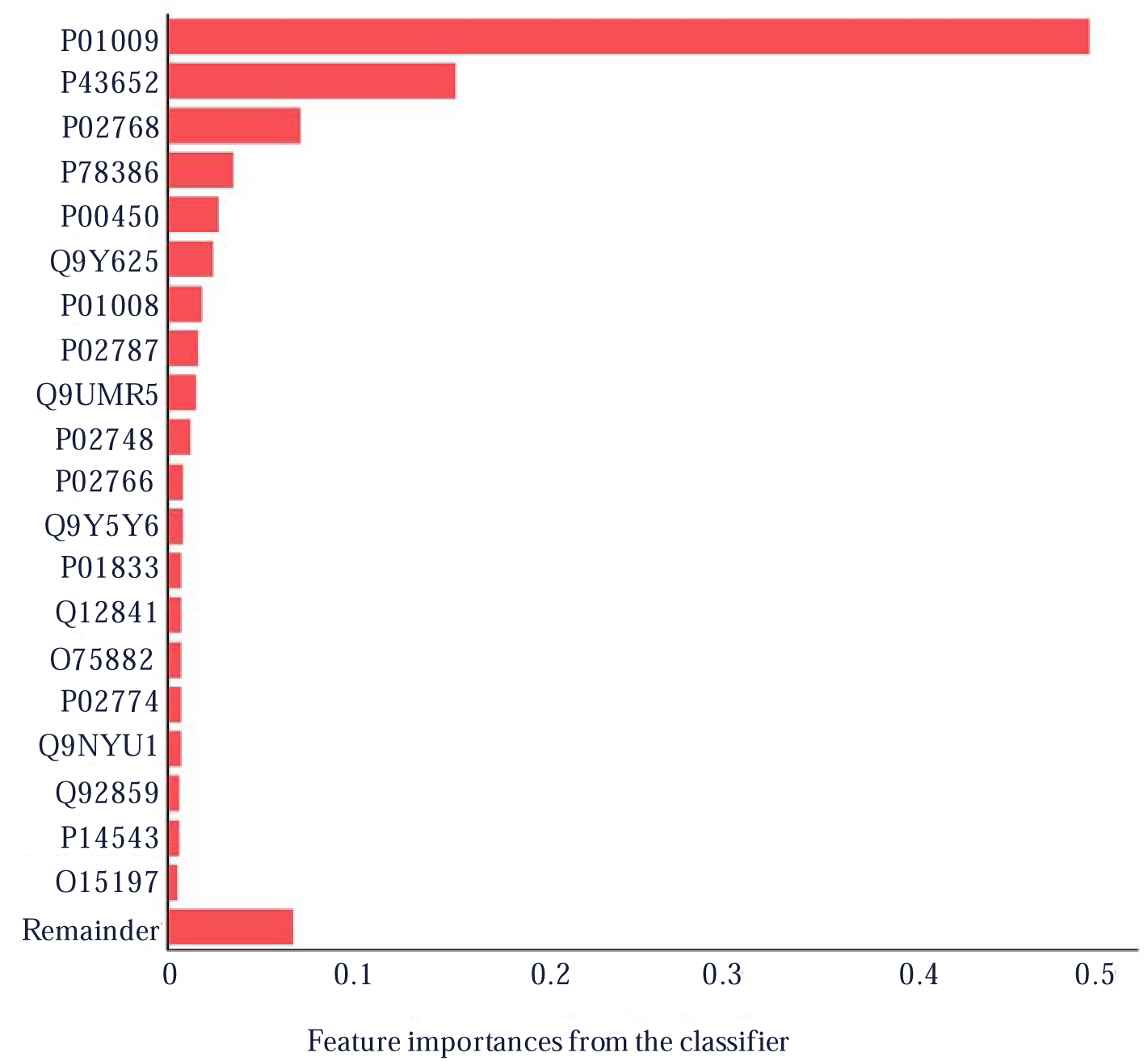
Top 20 features identified from the classifier using the XGBoost algorithm. UniProt IDs for each protein are shown.

### Functional annotation of proteins with significantly different abundances between cases and controls

Three hundred and thirty-two proteins with significantly different abundances between cases and controls were constructed into a network and annotated using Cytoscape. To gain more insight on the biological significance of these proteins, STRING enrichment was performed. The network was split into sub clusters based on relevant pathways related to CKD. We identified 112 proteins associated with the immune system (q-value [FDR]=1.4×10^-45^), and 89 proteins associated with the innate immune system (q=1.1×10^-32^) (**Supplementary Figures 4 and 5**). Additionally, 17 and six proteins were associated with extracellular matrix (ECM) organisation (q=0.03) and activation of matrix metalloproteinases (q=0.04), respectively (**Supplementary Figure 6A and B**).

Using Cytoscape public databases, 14 common proteins were identified between this pilot case-control study and other studies on CKD and hypertension, as imported from public databases (**Figure 6)**. The size of the circle represents the disease score, where a score of 0 indicates that the protein is not related to the disease and a score of 5 indicates that the protein is related to the disease. Proteins angiotensinogen (AGT), albumin (ALB), APOL1, and uromodulin (UMOD) were associated with CKD and hypertension with high disease scores of 5.0 (100% confidence), 4.0 (80%), 3.9 (78%), and 3.8 (76%), respectively. The proteins matrix metalloproteinase 9 (MMP9) and TIMP Metallopeptidase Inhibitor 1 (TIMP1) were associated with pathways involved in degradation of the ECM, interleukin (IL)-4 and IL-13 signalling, and activation of matrix metalloproteinases.

**Figure 6.**
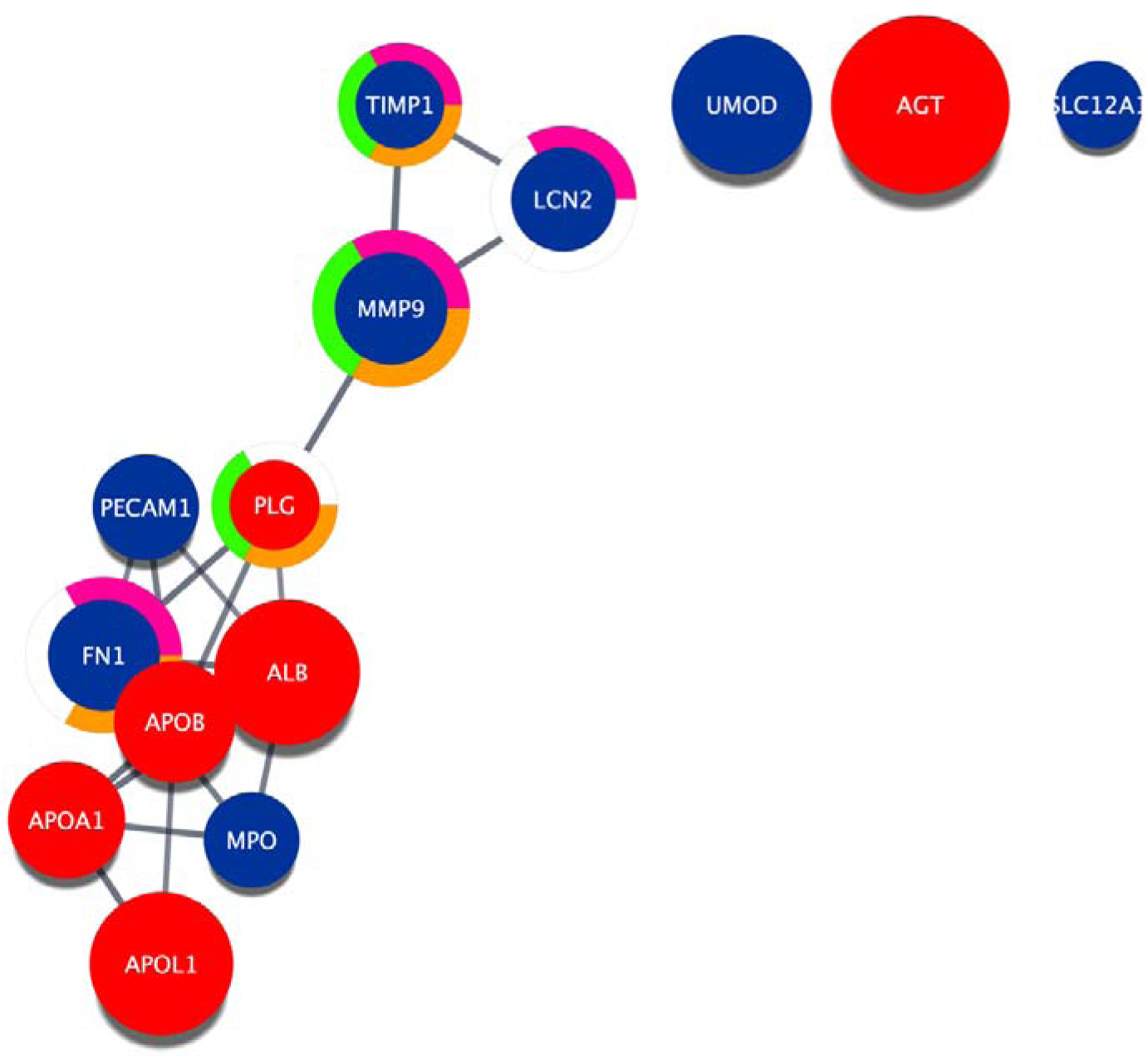
Common proteins between this case-control pilot study and other studies on CKD and hypertension, as imported from public databases. The gene names for each protein in the network are shown. Red indicates proteins with higher abundance in cases while blue represents proteins with lower abundance. Size of the circle represents the disease score (0 [not related to disease] – 5 [related to disease]). The closer to 5, the higher the confidence that the protein is associated with CKD and hypertension. AGT=5.0; ALB=4.0; APOL1=3.9; UMOD=3.8; APOB=3.2; APOA1=3.1; MMP9=2.9; FN1=2.9; PECAM1=2.7; LCN2=2.7; MPO=2.4; PLG=2.2; TIMP1=2.2; SLC12A1=2.2. Pathways are annotated on the network using a split donut ring. Orange indicates proteins associated with degradation of the extracellular matrix; pink indicates proteins associated with IL-4 and IL-13 signalling; green indicates proteins associated with activation of matrix metalloproteinases. CKD, chronic kidney disease; IL, interleukin.

## Discussion

This case-control pilot study aimed to identify potential proteins and pathways involved in hypertension-associated albuminuria by assessing urinary proteomic profiles in black South African participants with both hypertension and albuminuria compared to those who have neither condition.

Peptide and protein CVs showed that the methods were reproducible and that the system and workflow contributed low technical variability. The instrumental drift observed throughout the data acquisition period was minimal, as shown in the commercial Hela digest system suitability data that were acquired from a fresh sample on each day. A larger drift was observed in the study-specific peptide pool system suitability data; however, this can be attributed to autosampler-associated degradation as the sample was kept in the autosampler throughout the data acquisition process. This suggests that most of the variation observed in the study can be attributed to biological variation. On average, more proteins (1,225) were quantified in the control group than in the case group (915). Possible explanations include that a minor difference in the missed cleavage rate (∼3%) was observed between the two groups and several high abundance proteins, such as ALB, *APOL1*, apolipoprotein A1 (*APOA1*), and apolipoprotein B (*APOB*) are significantly higher in abundance in the cases ‘masking’ identification of lower abundance proteins, resulting in fewer quantifications overall.

The majority of differentially abundant proteins were associated with pathways involving the immune system, innate immune system, extracellular matrix organisation and activation of matrix metalloproteinases. In the setting of sustained hypertension, high blood pressure can cause permanent damage to the nephrons through arterial injury and glomerular ischaemia (40). Damage from renal ischaemia can lead to the production of various inflammatory cytokines that lead to immune cell infiltration and tubular atrophy (40). The latter is supported by Mattson (2014), (41) who showed an infiltration of inflammatory mononuclear cells in the arterioles and small arteries in kidney tissue from patients with hypertension. Enrichment analysis has shown that pathways related to inflammation and immune response are associated with the progression of kidney disease (42).

We identified six proteins associated with the activation of matrix metalloproteinase (MMPs) pathways. Of these, five had a lower abundance in participants with hypertension-associated albuminuria. It has been suggested that decreased activity of matrix metalloproteases may cause an accumulation of proteins in the extracellular matrix leading to fibrosis, which is one of the features of hypertension-associated kidney disease (43–45). Most of the proteins (71%; n=12/17) associated with the ECM organisation pathway were also reduced in participants with hypertension-associated albuminuria. It has been suggested that the reduced abundance of ECM organisation proteins may reflect reduced ECM turnover (decreased degradation), subsequently causing an increased deposit in ECM, resulting in fibrosis (46). Currently, kidney biopsy is the only validated approach to evaluate fibrosis. The identification of ECM organisation proteins may provide a non-invasive approach to detect early modifications in the ECM that lead to renal fibrosis.

Fourteen common proteins were identified between this pilot study and public databases on CKD and hypertension. Proteins including AGT, ALB, APOL1, and UMOD had the highest disease scores (76–100% confidence) for CKD and hypertension. In this pilot study, UMOD had a lower abundance in cases with hypertension-associated albuminuria. Similarly, Nqebelele et al. 2019 found reduced levels of UMOD in black African individuals with hypertensive-attributed CKD (47). Navise et al. (2023) also found lower levels of UMOD in black African individuals with CKD (48). In addition to its role in ECM remodelling and fibrosis, UMOD has been involved in regulating ion transport in the kidney (49). Reduced levels of UMOD have been postulated to be a result of decreased secretion from damaged tubules (50). Results from a mouse model with altered UMOD expression have shown that the induction of pro-inflammatory signalling is one of the first events that occur in the kidneys (42). The animal study also showed that the initiation of inflammatory signals precedes fibrosis and kidney damage, and possibly plays a vital role in disease onset (42).

It is important to note that hypercholesterolaemia was present in 38% of the cases in this study. Several studies have linked apolipoproteins (*APOA1*, *APOB* and *APOL1*) with CKD (51–53) and hypertension (54). Moreover, a Mendelian-randomization study identified a causal association between lipids (including low-density lipoprotein and triglycerides) and kidney function in individuals of African ancestry (55). It has been postulated that the association between apolipoproteins and kidney disease may be mediated by the impact of these lipoprotein particles on the kidney. Apolipoproteins and their associated lipids may have toxic effects on glomerular cells leading to glomerulosclerosis, which is scarring of the glomerulus (56). As a result, Kidney Disease: Improving Global Outcomes (KDIGO) developed guidelines for lipid management in patients with CKD (57).

Machine learning enabled us to train a classifier that on average correctly identified cases with an 88% true positive rate. Additionally, it identified controls with a 97% true negative rate. This analysis revealed that alpha-1-antitrypsin (P01009) and afamin (P43652) were the two most important features of the classifier. Studies have shown that alpha-1-antitrypsin and afamin are potential biomarkers for the diagnosis of early diabetic kidney disease (DKD) and can predict the decline in renal function (58, 59). Interestingly, in this pilot study, we have identified DKD markers in normoglycemia participants who have hypertension and albuminuria, which suggests the generalisability of the findings.

This is the first study to characterise the urinary proteome in South African individuals with hypertension-associated albuminuria. This case-control study identified common proteins with previous studies on CKD and hypertension from other ethnicities, and proteins associated with DKD, indicating generalisability of the findings to other populations and diseases. This study contributes to the understanding of pathways involved in hypertension-associated albuminuria and suggest opportunities for translation into the clinical setting.

The results from this study must be interpreted in context of its limitations, which included a relatively small sample size, consisting of only 73 individuals. All participants are from a single centre, therefore, results may not be generalisable.

Further research is essential to validate these outcomes in larger cohorts. In addition, the investigation of other diseases such as DKD and acute kidney injury would help to further explore generalisability of findings.

## Conclusions

In this pilot study, pathways associated with hypertension-associated albuminuria included the immune system, innate immune system, ECM organisation and activation of matrix metalloproteinases. These pathways contribute insights into the pathophysiology (e.g., immune cell infiltration, tubular atrophy, and ECM remodelling) of hypertension-related albuminuria. Additionally, proteins such as AGT, ALB, *APOL1* and UMOD had the highest disease scores (76–100% confidence) for hypertension and CKD. The urinary proteomic data combined with the machine learning approach classified disease status and identified proteins and pathways associated with hypertension and albuminuria.

## List of abbreviations

ARK: African Research Kidney Disease
CKD: chronic kidney disease
DB: diastolic blood pressure
DIA: data-independent acquisition
ECM: extracellular matrix
FDR: false discovery rate
iRT: normalised retention time
LCMS: liquid chromatography with mass spectrometry
MeCN: acetonitrile
PCA: principal component analysis
QC: quality control
RT: room temperature
SBP: systolic blood pressure
SWATH: Sequential Window Acquisition of all Theoretical Mass Spectra
UHPLC: ultra-high-performance liquid chromatography

## Declarations

Ethics approval and consent to participate

The study was approved by the Human Research Ethics Committee (Medical) of the University of the Witwatersrand (certificate number, M200331). All participants provided written informed consent.

## Author contributions

All authors have contributed to the design of the study. MAG and ISG wrote the main manuscript text and prepared figures. ISG, SHS, MR, JF, and J-TB critically reviewed the manuscript.

## Consent for publication

This manuscript has been read and approved by all the authors to publish and is not submitted or under consideration for publication elsewhere.

## Data availability

The proteomic data files described in this study have been deposited to the ProteomeXchange Consortium via the PRIDE partner repository with the dataset identifier (PXD046441) (60).

## Competing interests

ISG and SHS are employees of ReSyn Biosciences who are the proprietors of MagReSyn® HILIC technology. The remaining authors have no conflicts of interest associated with this manuscript.

## References

1. World Health Organization. Hypertension. Available at: https://www.who.int/news-room/fact-sheets/detail/hypertension. (accessed August 2023).

2. Hariparshad S, Bhimma R, Nandlal L, Jembere E, Naicker S, Assounga A. The prevalence of chronic kidney disease in South Africa-limitations of studies comparing prevalence with sub-Saharan Africa, Africa, and globally. BMC nephrology. 2023;24(1):62.

3. De Bhailis ÁM, Kalra PA. Hypertension and the kidneys. British Journal of Hospital Medicine. 2022;83(5):1–11.

4. Zandi-Nejad K, Luyckx VA, Brenner BM. Adult hypertension and kidney disease: the role of fetal programming. Hypertension. 2006;47(3):502–8.

5. Gjerde A. Low birth weight, intrauterine growth restriction and risk of chronic kidney disease in adult age. 2022.

6. Patel P, Sanghavi DK, Morris DL, et al. Angiotensin II. [Updated 2023 May 26]. In: StatPearls [Internet]. Treasure Island (FL): StatPearls Publishing; 2023 Jan-. Available from: https://www.ncbi.nlm.nih.gov/books/NBK499912/. (accessed August 2023).

7. Benigni A, Cassis P, Remuzzi G. Angiotensin II revisited: new roles in inflammation, immunology and aging. EMBO molecular medicine. 2010;2(7):247–57.

8. Pollock DM. Renal endothelin in hypertension. Current opinion in nephrology and hypertension. 2000;9(2):157–64.

9. Sasser JM, Pollock JS, Pollock DM. Renal endothelin in chronic angiotensin II hypertension. American Journal of Physiology-Regulatory, Integrative and Comparative Physiology. 2002;283(1):R243–R8.

10. Navar L, Inscho E, Majid S, Imig J, Harrison-Bernard L, Mitchell K. Paracrine regulation of the renal microcirculation. Physiological reviews. 1996;76(2):425–536.

11. Rüster C, Wolf G. Renin-angiotensin-aldosterone system and progression of renal disease. Journal of the American Society of Nephrology. 2006;17(11):2985–91.

12. Siragy HM, Carey RM. Role of the intrarenal renin-angiotensin-aldosterone system in chronic kidney disease. American journal of nephrology. 2010;31(6):541–50.

13. Bidani AK, Griffin KA. Pathophysiology of hypertensive renal damage: implications for therapy. Hypertension. 2004;44(5):595–601.

14. Shankland S. The podocyte’s response to injury: role in proteinuria and glomerulosclerosis. Kidney international. 2006;69(12):2131–47.

15. Folkow B, Göthberg G, Lundin S, Ricksten SE. Structural “resetting” of the renal vascular bed in spontaneously hypertensive rats (SHR). Acta physiologica Scandinavica. 1977;100(2):270–2.

16. Buffet L, Ricchetti C. Chronic kidney disease and hypertension: A destructive combination. 2012.

17. Heerspink HJL, Gansevoort RT. Albuminuria is an appropriate therapeutic target in patients with CKD: the pro view. Clinical Journal of the American Society of Nephrology. 2015;10(6):1079–88.

18. Levin A, Stevens PE, Bilous RW, Coresh J, De Francisco AL, De Jong PE, et al. Kidney Disease: Improving Global Outcomes (KDIGO) CKD Work Group. KDIGO 2012 clinical practice guideline for the evaluation and management of chronic kidney disease. Kidney international supplements. 2013;3(1):1–150.

19. Lopez-Giacoman S, Madero M. Biomarkers in chronic kidney disease, from kidney function to kidney damage. World journal of nephrology. 2015;4(1):57.

20. Heerspink HJL, Gansevoort RT. Albuminuria is an appropriate therapeutic target in patients with CKD: the pro view. Clinical journal of the American Society of Nephrology: CJASN. 2015;10(6):1079.

21. Good DM, Zürbig P, Argiles A, Bauer HW, Behrens G, Coon JJ, et al. Naturally occurring human urinary peptides for use in diagnosis of chronic kidney disease. Molecular & cellular proteomics. 2010;9(11):2424–37.

22. Wu I-W, Tsai T-H, Lo C-J, Chou Y-J, Yeh C-H, Chan Y-H, et al. Discovering a trans-omics biomarker signature that predisposes high risk diabetic patients to diabetic kidney disease. npj Digital Medicine. 2022;5(1):166.

23. Cisek K, Krochmal M, Klein J, Mischak H. The application of multi-omics and systems biology to identify therapeutic targets in chronic kidney disease. Nephrology Dialysis Transplantation. 2016;31(12):2003–11.

24. Provenzano M, Serra R, Garofalo C, Michael A, Crugliano G, Battaglia Y, et al. OMICS in Chronic Kidney Disease: Focus on Prognosis and Prediction. International Journal of Molecular Sciences. 2021;23(1):336.

25. Mischak H, Delles C, Klein J, Schanstra JP. Urinary proteomics based on capillary electrophoresis-coupled mass spectrometry in kidney disease: discovery and validation of biomarkers, and clinical application. Advances in chronic kidney disease. 2010;17(6):493–506.

26. Fan G, Gong T, Lin Y, Wang J, Sun L, Wei H, et al. Urine proteomics identifies biomarkers for diabetic kidney disease at different stages. Clinical Proteomics. 2021;18(1):1–12.

27. Pontillo C, Mischak H. Urinary peptide-based classifier CKD273: towards clinical application in chronic kidney disease. Clinical kidney journal. 2017;10(2):192–201.

28. De Beer D, Mels CM, Schutte AE, Delles C, Mary S, Mullen W, et al. Identifying a urinary peptidomics profile for hypertension in young adults: The AfricanCPREDICT study: Urinary peptidomics and hypertension. Proteomics. 2023;23(11):2200444.

29. Kalyesubula R, Fabian J, Nakanga W, Newton R, Ssebunnya B, Prynn J, et al. How to estimate glomerular filtration rate in sub-Saharan Africa: design and methods of the African Research into Kidney Diseases (ARK) study. BMC nephrology. 2020;21(1):1–12.

30. Craik A, Gondwe M, Mayindi N, Chipungu S, Khoza B, Gómez-Olivé X, et al. Forgotten but not gone in rural South Africa: Urinary schistosomiasis and implications for chronic kidney disease screening in endemic countries [version 1; peer review: awaiting peer review]. 2023.

31. Craik A, Gondwe M, Mayindi N, Chipungu S, Khoza B, Gómez-Olivé X, et al. Forgotten but not gone in rural South Africa: Urinary schistosomiasis and implications for chronic kidney disease screening in endemic countries. Wellcome Open Research. 2023;8(68):68.

32. George JA, Brandenburg J-T, Fabian J, Crowther NJ, Agongo G, Alberts M, et al. Kidney damage and associated risk factors in rural and urban sub-Saharan Africa (AWI-Gen): a cross-sectional population study. The Lancet Global Health. 2019;7(12):e1632–e43.

33. Nweke EE, Naicker P, Aron S, Stoychev S, Devar J, Tabb DL, et al. SWATH-MS based proteomic profiling of pancreatic ductal adenocarcinoma tumours reveals the interplay between the extracellular matrix and related intracellular pathways. Plos one. 2020;15(10):e0240453.

34. Govender IS, Mokoena R, Stoychev S, Naicker P. Urine-HILIC: Automated sample preparation for bottom-up urinary proteome profiling in clinical proteomics. Proteomes. 2023;11(4):29.

35. Röst HL, Rosenberger G, Navarro P, Gillet L, Miladinović SM, Schubert OT, et al. OpenSWATH enables automated, targeted analysis of data-independent acquisition MS data. Nature biotechnology. 2014;32(3):219–23.

36. Reiter L, Rinner O, Picotti P, Hüttenhain R, Beck M, Brusniak M-Y, et al. mProphet: automated data processing and statistical validation for large-scale SRM experiments. Nature methods. 2011;8(5):430–5.

37. Zhang B, Chambers MC, Tabb DL. Proteomic parsimony through bipartite graph analysis improves accuracy and transparency. Journal of proteome research. 2007;6(9):3549–57.

38. Shannon P, Markiel A, Ozier O, Baliga NS, Wang JT, Ramage D, et al. Cytoscape: a software environment for integrated models of biomolecular interaction networks. Genome research. 2003;13(11):2498–504.

39. Torun FM, Virreira Winter S, Doll S, Riese FM, Vorobyev A, Mueller-Reif JB, et al. Transparent exploration of machine learning for biomarker discovery from proteomics and omics data. Journal of Proteome Research. 2022;22(2):359–67.

40. Benson LN, Guo Y, Deck K, Mora C, Liu Y, Mu S. The link between immunity and hypertension in the kidney and heart. Frontiers in Cardiovascular Medicine. 2023;10.

41. Mattson DL. Infiltrating immune cells in the kidney in salt-sensitive hypertension and renal injury. American Journal of Physiology-Renal Physiology. 2014;307(5):F499–F508.

42. Trudu M, Schaeffer C, Riba M, Ikehata M, Brambilla P, Messa P, et al. Early involvement of cellular stress and inflammatory signals in the pathogenesis of tubulointerstitial kidney disease due to UMOD mutations. Scientific reports. 2017;7(1):7383.

43. Hughson MD, Puelles VG, Hoy WE, Douglas-Denton RN, Mott SA, Bertram JF. Hypertension, glomerular hypertrophy and nephrosclerosis: the effect of race. Nephrology Dialysis Transplantation. 2014;29(7):1399–409.

44. Bazzi C, Seccia TM, Napodano P, Campi C, Caroccia B, Cattarin L, et al. High blood pressure is associated with tubulointerstitial damage along with glomerular damage in glomerulonephritis. A large cohort study. Journal of Clinical Medicine. 2020;9(6):1656.

45. Cheng S, Pollock AS, Mahimkar R, Olson JL, Lovett DH, Cheng S, et al. Matrix metalloproteinase 2 and basement membrane integrity: a unifying mechanism for progressive renal injury. The FASEB journal. 2006;20(11):1898–900.

46. Nkuipou-Kenfack E, Duranton F, Gayrard N, Argiles A, Lundin U, Weinberger KM, et al. Assessment of metabolomic and proteomic biomarkers in detection and prognosis of progression of renal function in chronic kidney disease. PloS one. 2014;9(5):e96955.

47. Nqebelele NU, Dickens C, Dix-Peek T, Duarte R, Naicker S. Urinary uromodulin levels and UMOD variants in black South Africans with hypertension-attributed chronic kidney disease. International Journal of Nephrology. 2019;2019.

48. Navise NH, Mokwatsi GG, Gafane-Matemane LF, Fabian J, Lammertyn L. Kidney dysfunction: prevalence and associated risk factors in a community-based study from the North West Province of South Africa. BMC nephrology. 2023;24(1):1–8.

49. Renigunta A, Renigunta V, Saritas T, Decher N, Mutig K, Waldegger S. Tamm-Horsfall glycoprotein interacts with renal outer medullary potassium channel ROMK2 and regulates its function. Journal of Biological Chemistry. 2011;286(3):2224–35.

50. Prajczer S, Heidenreich U, Pfaller W, Kotanko P, Lhotta K, Jennings P. Evidence for a role of uromodulin in chronic kidney disease progression. Nephrology Dialysis Transplantation. 2010;25(6):1896–903.

51. Goek O-N, Köttgen A, Hoogeveen RC, Ballantyne CM, Coresh J, Astor BC. Association of apolipoprotein A1 and B with kidney function and chronic kidney disease in two multiethnic population samples. Nephrology Dialysis Transplantation. 2012;27(7):2839–47.

52. Zhao W-b, Alberto DLPSM. Serum apolipoprotein B/apolipoprotein A1 ratio is associated with the progression of diabetic kidney disease to renal replacement therapy. International Urology and Nephrology. 2020;52:1923–8.

53. Ma L, Divers J, Freedman BI. Mechanisms of injury in APOL1-associated kidney disease. Transplantation. 2019;103(3):487.

54. Nayak P, Panda S, Thatoi PK, Rattan R, Mohapatra S, Mishra PK. Evaluation of lipid profile and apolipoproteins in essential hypertensive patients. Journal of Clinical and Diagnostic Research: JCDR. 2016;10(10):BC01.

55. Kintu C, Soremekun O, Kamiza AB, Kalungi A, Mayanja R, Kalyesubula R, et al. The causal effects of lipid traits on kidney function in Africans: bidirectional and multivariable Mendelian-randomization study. EBioMedicine. 2023;90.

56. Kwon S, Kim DK, Oh K-H, Joo KW, Lim CS, Kim YS, et al. Apolipoprotein B is a risk factor for end-stage renal disease. Clinical Kidney Journal. 2021;14(2):617–23.

57. Wanner C, Tonelli M. KDIGO Clinical Practice Guideline for Lipid Management in CKD: summary of recommendation statements and clinical approach to the patient. Kidney international. 2014;85(6):1303–9.

58. Ning J, Xiang Z, Xiong C, Zhou Q, Wang X, Zou H. Alpha1-antitrypsin in urinary extracellular vesicles: A potential biomarker of diabetic kidney disease prior to microalbuminuria. Diabetes, Metabolic Syndrome and Obesity. 2020:2037–48.

59. Kaburagi Y, Takahashi E, Kajio H, Yamashita S, Yamamoto-Honda R, Shiga T, et al. Urinary afamin levels are associated with the progression of diabetic nephropathy. Diabetes research and clinical practice. 2019;147:37–46.

60. Perez-Riverol Y, Bai J, Bandla C, García-Seisdedos D, Hewapathirana S, Kamatchinathan S, et al. The PRIDE database resources in 2022: a hub for mass spectrometry-based proteomics evidences. Nucleic acids research. 2022;50(D1):D543–D52.

