## Supplementary data for "Proteomic insights into the pathophysiology of hypertension-associated albuminuria: Pilot study in a South African cohort"

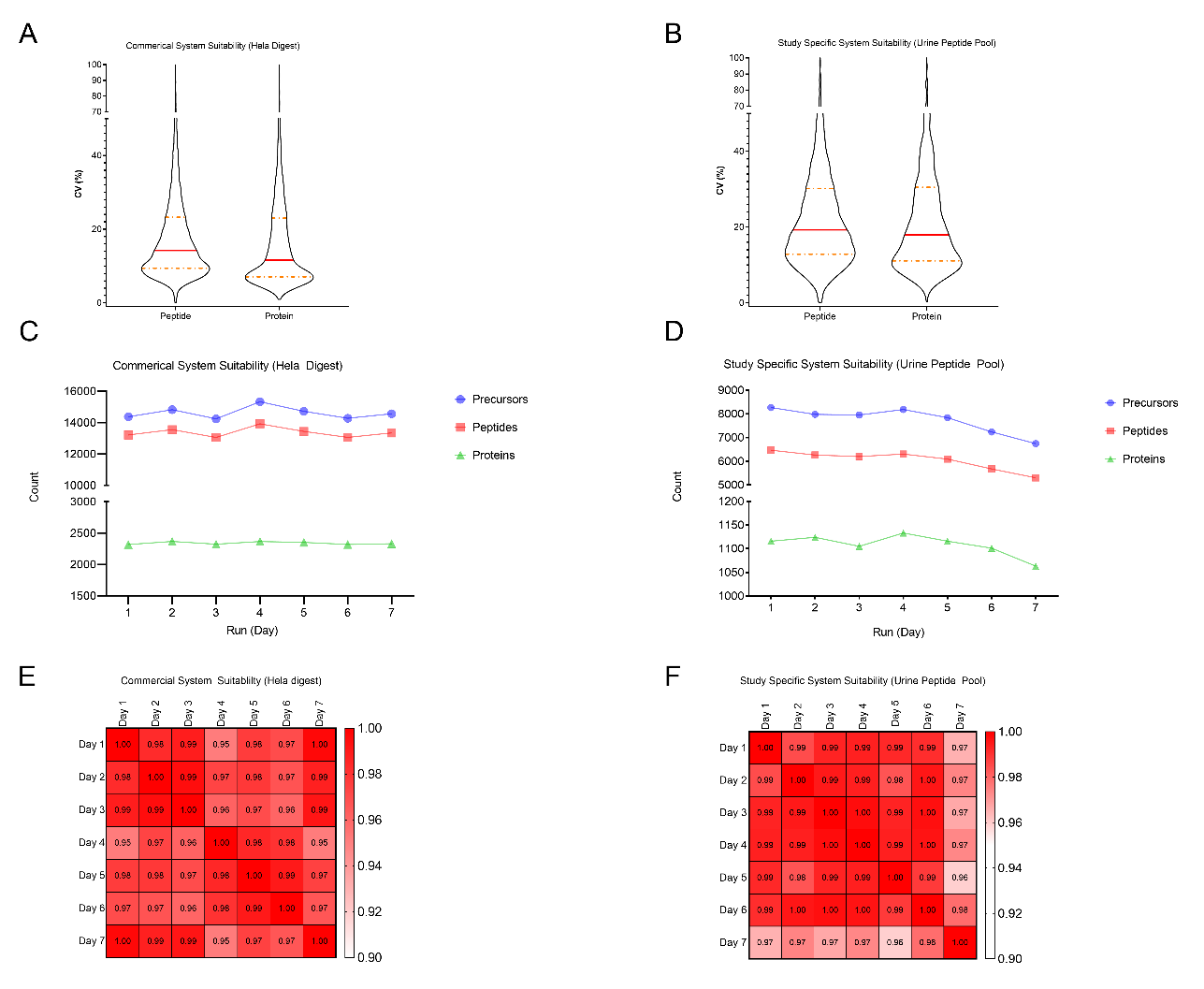

**Supplementary Figure 1. Project specific system suitability-quality control.**

CV, co-efficient of variation.

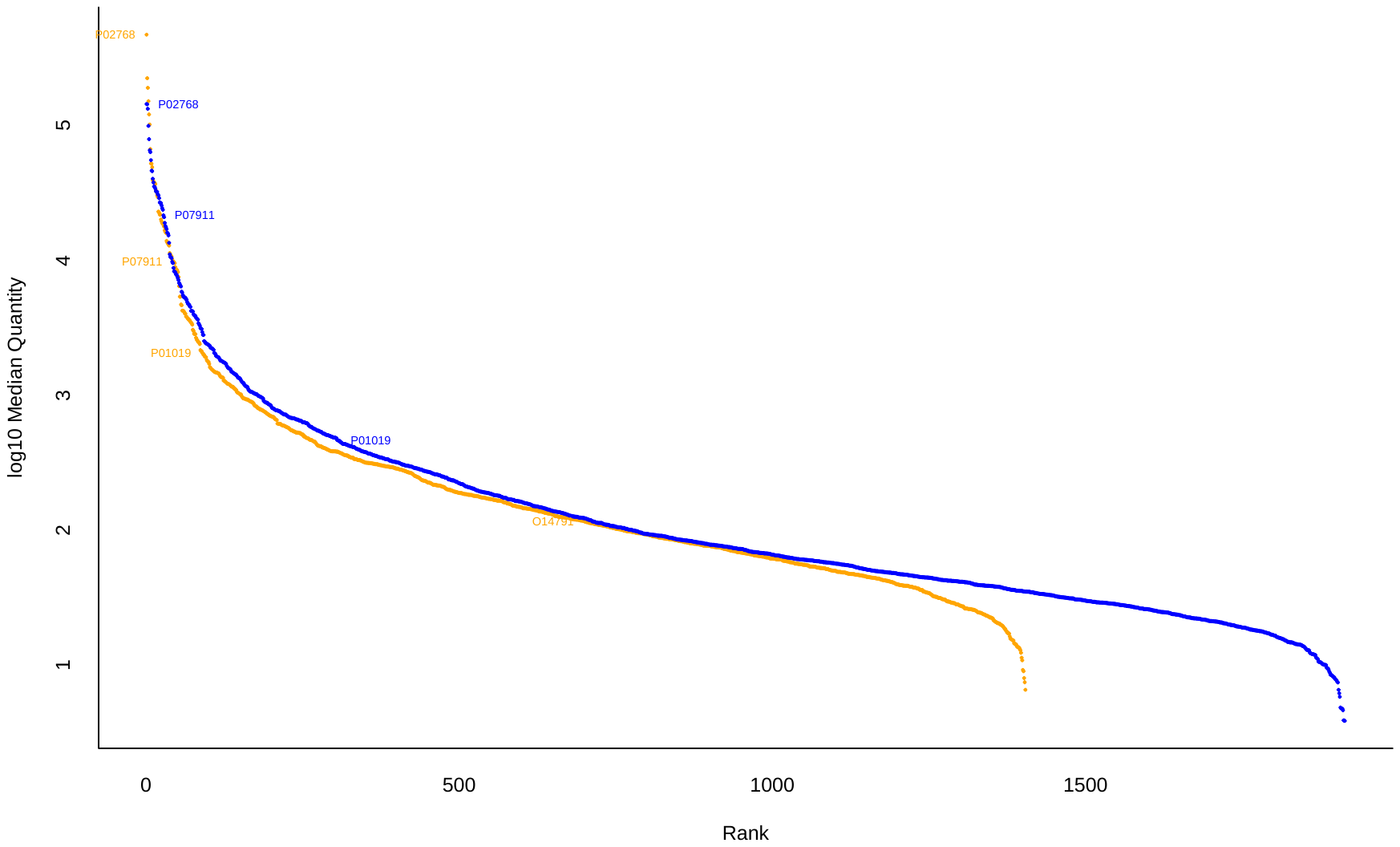

**Supplementary Figure 2. Dynamic range of proteins quantified.**

Orange dots indicate cases and blue dots indicate controls. The high disease scoring proteins are annotated.

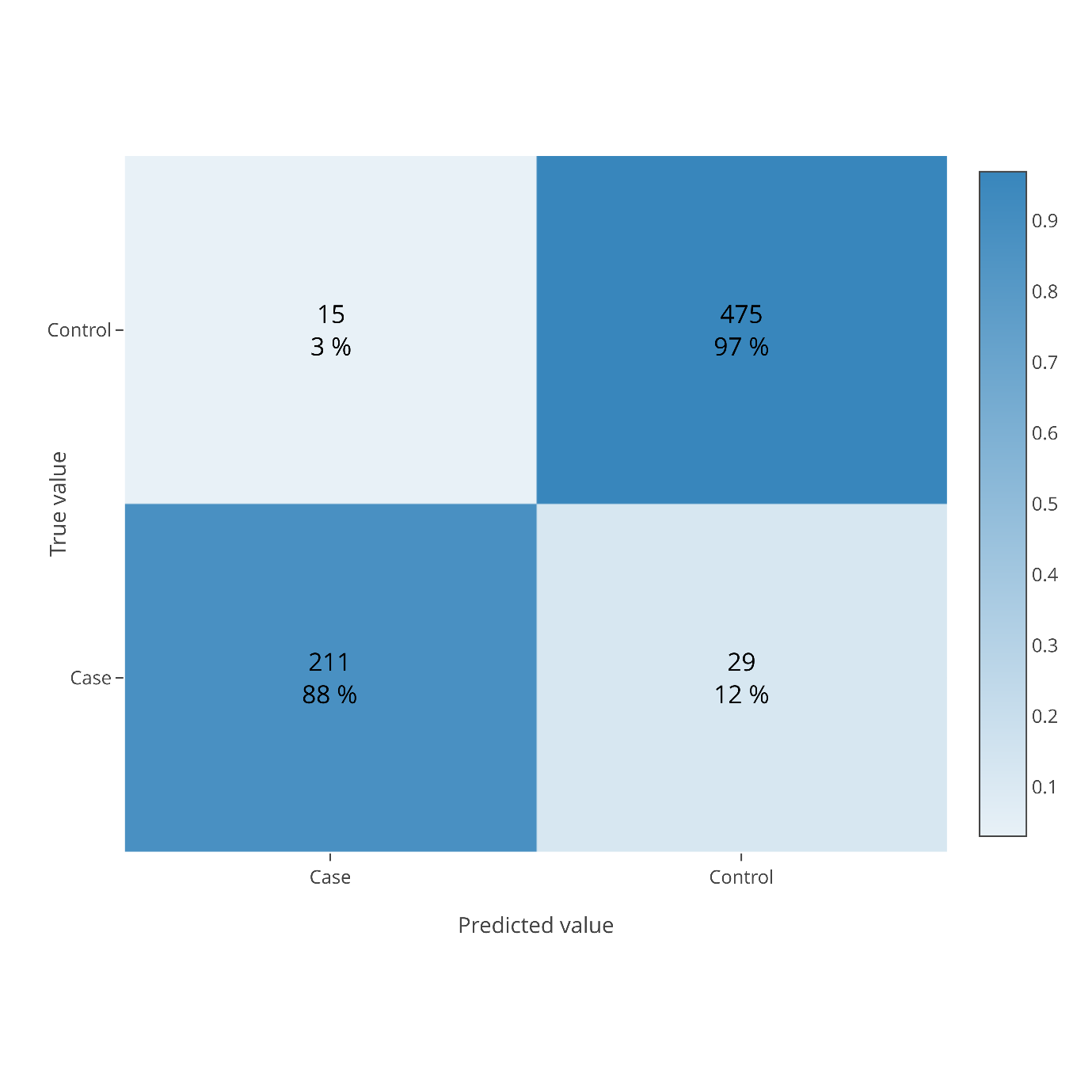

**Supplementary Figure 3: Confusion matrix following machine learning using the XGBoost algorithm.**

Confusion matrix refers to the percentage of samples assigned to each group.

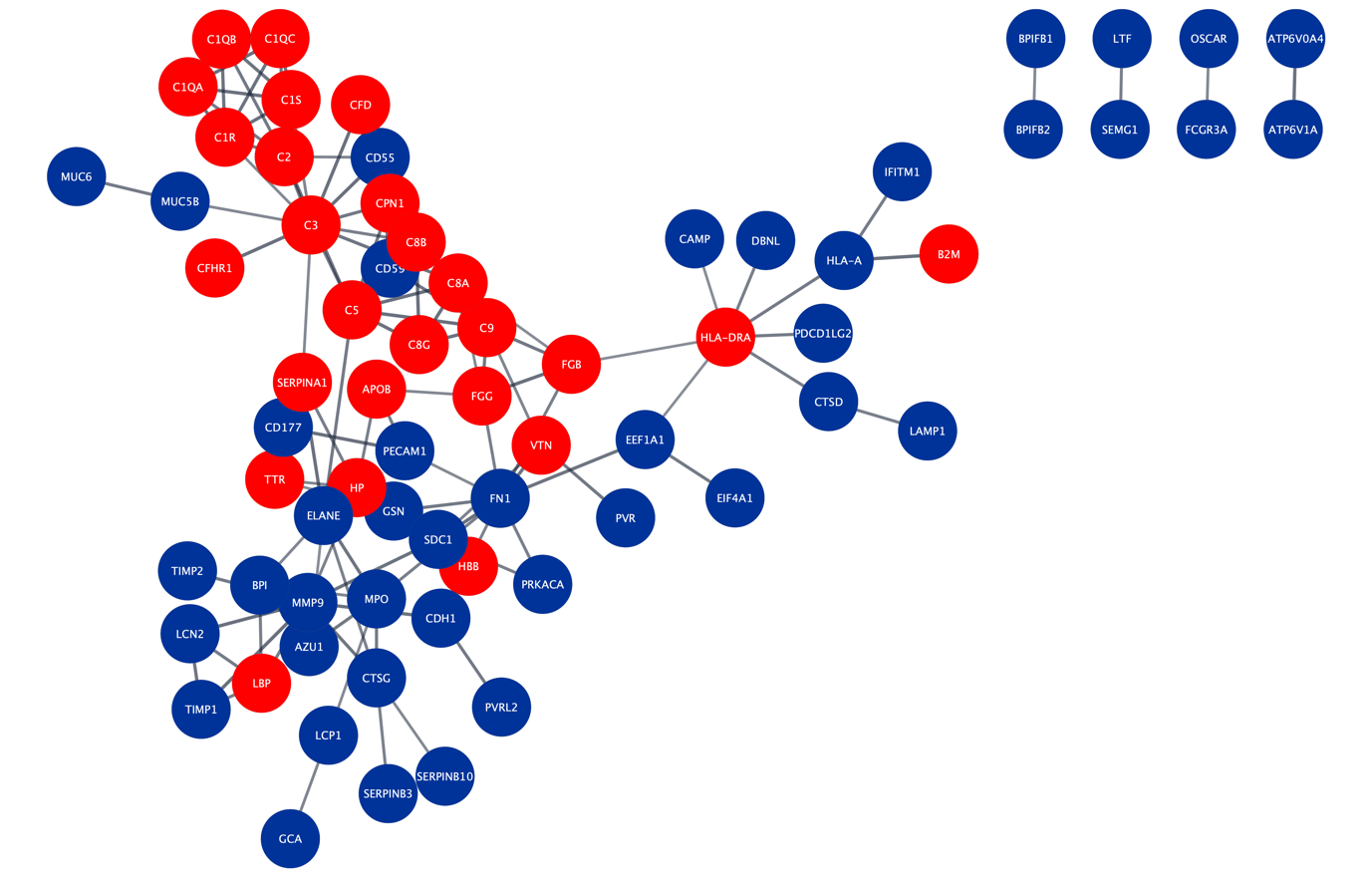

**Supplementary Figure 4. Differentially abundant proteins associated with the immune system.**

The gene names for each protein in the network are shown. Differentially abundant proteins were detected at an FDR of 0.01 and a fold-change of 2. Red indicates proteins with higher abundance in cases while blue represents proteins with lower abundance.

FDR, false discovery rate.

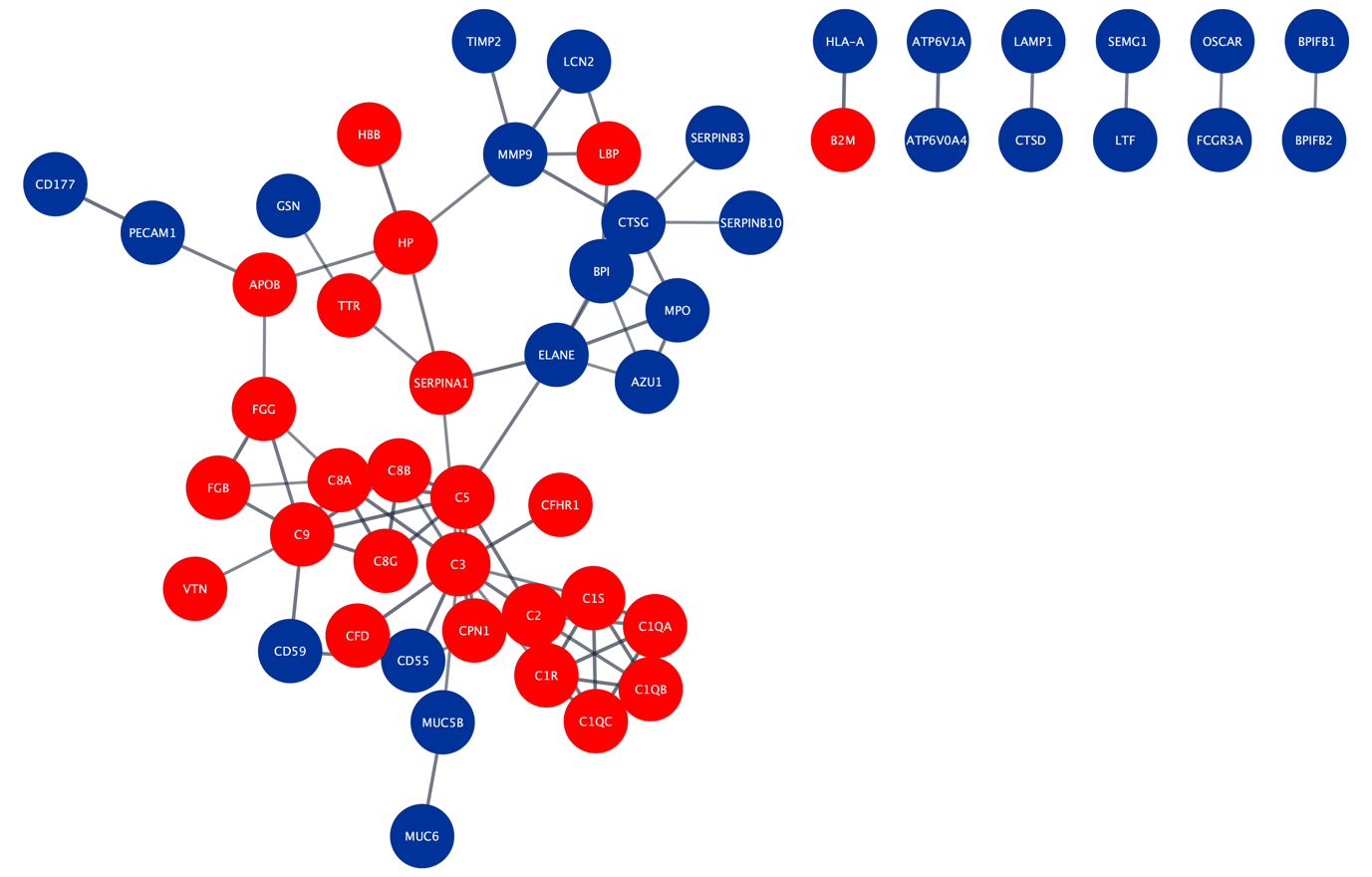

**Supplementary Figure 5. Differentially abundant proteins associated with the innate immune system.**

The gene names for each protein in the network are shown. Differentially abundant proteins were detected at an FDR of 0.01 and a fold-change of 2. Red indicates proteins with higher abundance in cases while blue represents proteins with lower abundance.

FDR, false discovery rate.

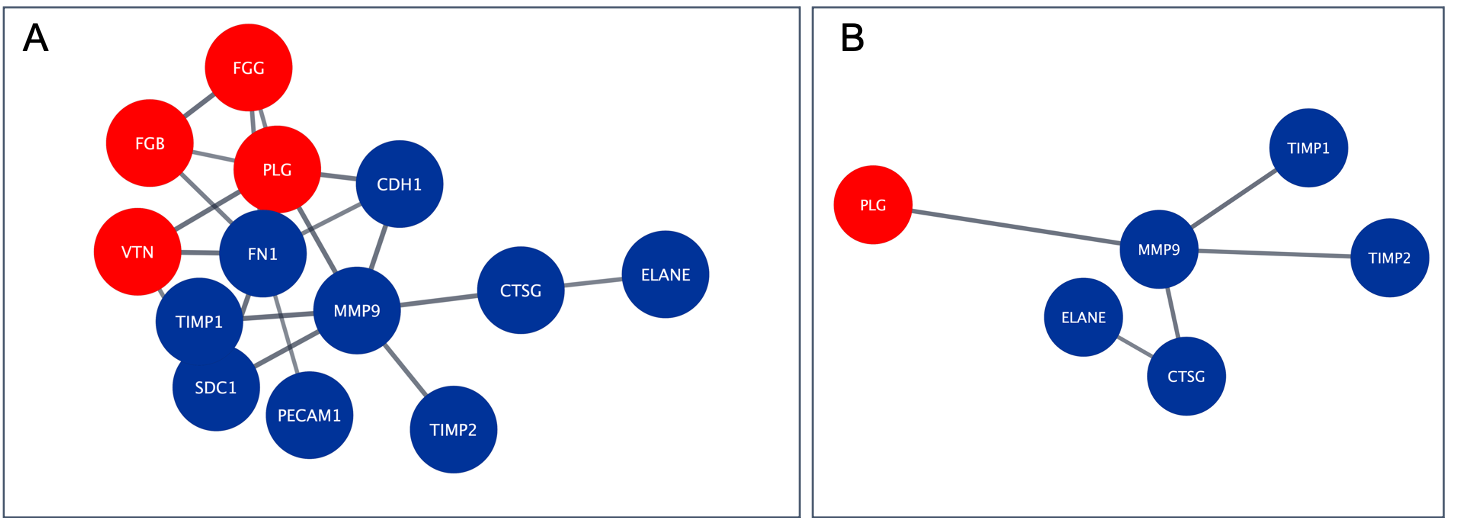

**Supplementary Figure 6. Differentially abundant proteins associated with extracellular matrix organisation (A) and activation of matrix metalloproteinases (B).**

The gene names for each protein in the network are shown. Differentially abundant proteins were detected at an FDR of 0.01 and a fold-change of 2. Red indicates proteins with higher abundance in cases while blue represents proteins with lower abundance.

FDR, false discovery rate.

**Supplementary Table 1. Significantly different proteins in the volcano plot**

| Protein Groups | **Difference** | **Direction** | **Log q-value** |
| --- | --- | --- | --- |
| P43652 | 4,01 | Positive | 14,79 |
| P25691; P78386 | 3,65 | Positive | 10,24 |
| Q9Y625 | 3,03 | Positive | 9,85 |
| P02787 | 3,00 | Positive | 13,76 |
| P02144 | 2,87 | Positive | 8,61 |
| P01024 | 2,58 | Positive | 7,00 |
| P01009 | 2,66 | Positive | 18,70 |
| P02647 | 2,51 | Positive | 5,45 |
| Q8WYP5 | 2,49 | Positive | 7,59 |
| P02766 | 2,43 | Positive | 7,54 |
| P01008 | 2,38 | Positive | 10,34 |
| P19823 | 2,35 | Positive | 5,40 |
| Q9UGM5 | 2,23 | Positive | 6,23 |
| P69905 | 2,15 | Positive | 2,55 |
| Q9NYQ6 | 2,10 | Positive | 2,80 |
| P06727 | 2,07 | Positive | 4,21 |
| P00450 | 2,00 | Positive | 8,21 |
| P02774 | 1,99 | Positive | 9,26 |
| P02748 | 1,99 | Positive | 7,57 |
| P02768 | 1,98 | Positive | 10,99 |
| P00747 | 1,93 | Positive | 5,55 |
| P36955 | 1,89 | Positive | 7,18 |
| P27169 | 1,85 | Positive | 4,34 |
| P00738 | 1,81 | Positive | 2,89 |
| P01019 | 1,80 | Positive | 5,35 |
| P01860 | 1,77 | Positive | 3,08 |
| P08697 | 1,77 | Positive | 7,77 |
| P06681 | 1,76 | Positive | 6,70 |
| P00739 | 1,73 | Positive | 3,73 |
| P18428 | 1,66 | Positive | 4,77 |
| P04217 | 1,64 | Positive | 5,17 |
| Q96KN2 | 1,50 | Positive | 5,10 |
| P07357 | 1,48 | Positive | 5,68 |
| P07360 | 1,47 | Positive | 3,36 |
| A0A0C4DH38 | 1,46 | Positive | 3,17 |
| P08185 | 1,37 | Positive | 3,76 |
| P28072 | -1,15 | Negative | 4,44 |
| Q9UN73; Q9Y5H7 | -1,16 | Negative | 4,15 |
| P28070 | -1,17 | Negative | 4,09 |
| P06733 | -1,18 | Negative | 4,79 |
| P60174 | -1,21 | Negative | 5,01 |
| Q12907 | -1,21 | Negative | 4,45 |
| P04275 | -1,22 | Negative | 4,13 |
| P05362 | -1,22 | Negative | 3,88 |
| P27930 | -1,22 | Negative | 4,15 |
| Q8TER0 | -1,22 | Negative | 6,23 |
| Q9HAT2 | -1,22 | Negative | 3,79 |
| O00462 | -1,23 | Negative | 5,53 |
| P05067 | -1,24 | Negative | 4,84 |
| O75083 | -1,24 | Negative | 4,40 |
| P16083 | -1,26 | Negative | 3,82 |
| A0A0C4DH25 | -1,27 | Negative | 3,92 |
| Q99497 | -1,27 | Negative | 4,76 |
| O00182 | -1,28 | Negative | 3,70 |
| Q15113 | -1,29 | Negative | 4,94 |
| Q08257 | -1,29 | Negative | 4,93 |
| Q99536 | -1,29 | Negative | 3,69 |
| P21709 | -1,30 | Negative | 4,12 |
| P50395 | -1,30 | Negative | 4,63 |
| Q04760 | -1,31 | Negative | 3,57 |
| O00241 | -1,31 | Negative | 3,57 |
| P68104; Q5VTE0 | -1,31 | Negative | 4,58 |
| P40121 | -1,32 | Negative | 6,02 |
| P41222 | -1,32 | Negative | 3,49 |
| Q9P2B2 | -1,32 | Negative | 5,32 |
| Q16651 | -1,32 | Negative | 4,51 |
| Q14894 | -1,33 | Negative | 4,81 |
| Q96J84 | -1,34 | Negative | 5,58 |
| P51148 | -1,34 | Negative | 4,85 |
| Q14393 | -1,34 | Negative | 4,11 |
| O94919 | -1,34 | Negative | 3,39 |
| P11021 | -1,34 | Negative | 4,88 |
| Q96JQ0 | -1,34 | Negative | 4,58 |
| P09619 | -1,35 | Negative | 3,77 |
| Q15223 | -1,35 | Negative | 4,84 |
| Q06830 | -1,36 | Negative | 5,40 |
| P07204 | -1,36 | Negative | 4,16 |
| P07711 | -1,37 | Negative | 6,11 |
| P13639 | -1,38 | Negative | 4,76 |
| P11233 | -1,39 | Negative | 4,37 |
| Q9H8L6 | -1,39 | Negative | 4,22 |
| P05090 | -1,39 | Negative | 3,94 |
| P38606 | -1,39 | Negative | 3,90 |
| Q96AP7 | -1,39 | Negative | 3,15 |
| Q86T13 | -1,40 | Negative | 5,49 |
| P23526 | -1,41 | Negative | 5,12 |
| Q13332 | -1,41 | Negative | 4,95 |
| P30086 | -1,41 | Negative | 4,22 |
| Q13621 | -1,41 | Negative | 3,45 |
| P23284 | -1,42 | Negative | 4,55 |
| P17174 | -1,42 | Negative | 4,20 |
| Q8WVQ1 | -1,43 | Negative | 6,00 |
| P61970 | -1,44 | Negative | 5,76 |
| P08582 | -1,45 | Negative | 5,80 |
| Q8N3J6 | -1,45 | Negative | 6,53 |
| P11597 | -1,45 | Negative | 3,13 |
| P14174 | -1,46 | Negative | 3,53 |
| P52758 | -1,46 | Negative | 3,49 |
| O95967 | -1,46 | Negative | 5,46 |
| P55285 | -1,47 | Negative | 4,18 |
| P35555 | -1,48 | Negative | 3,55 |
| Q7Z7M0 | -1,48 | Negative | 4,99 |
| Q14118 | -1,48 | Negative | 3,66 |
| Q9HCU0 | -1,49 | Negative | 4,99 |
| P04156 | -1,50 | Negative | 4,36 |
| Q9Y624 | -1,51 | Negative | 4,05 |
| O00757 | -1,51 | Negative | 2,98 |
| P05937 | -1,51 | Negative | 3,67 |
| Q16769 | -1,52 | Negative | 3,44 |
| P12109 | -1,52 | Negative | 5,28 |
| P15941 | -1,53 | Negative | 3,22 |
| O60888 | -1,53 | Negative | 3,06 |
| O00391 | -1,54 | Negative | 6,36 |
| P0DJD8 | -1,55 | Negative | 4,96 |
| Q6UX71 | -1,55 | Negative | 6,07 |
| P01133 | -1,55 | Negative | 4,05 |
| P12111 | -1,55 | Negative | 4,38 |
| P06396 | -1,56 | Negative | 3,64 |
| Q10588 | -1,56 | Negative | 4,48 |
| Q99983 | -1,56 | Negative | 4,85 |
| P42785 | -1,56 | Negative | 4,36 |
| P19022 | -1,57 | Negative | 3,82 |
| P14618 | -1,57 | Negative | 3,55 |
| O75882 | -1,57 | Negative | 5,56 |
| P13645 | -1,57 | Negative | 2,95 |
| A6NI73 | -1,57 | Negative | 5,77 |
| Q53RD9 | -1,57 | Negative | 4,44 |
| Q01459 | -1,58 | Negative | 4,02 |
| P29323 | -1,58 | Negative | 6,59 |
| Q92692 | -1,58 | Negative | 4,22 |
| P08572 | -1,58 | Negative | 3,66 |
| P15121 | -1,59 | Negative | 6,18 |
| P00966 | -1,59 | Negative | 2,82 |
| Q9HCN6 | -1,60 | Negative | 3,51 |
| O00115 | -1,60 | Negative | 6,66 |
| Q6UXB4 | -1,60 | Negative | 3,35 |
| O95865 | -1,61 | Negative | 3,90 |
| P98160 | -1,61 | Negative | 3,82 |
| Q12860 | -1,61 | Negative | 8,78 |
| Q8WW52 | -1,62 | Negative | 3,89 |
| P16035 | -1,62 | Negative | 8,09 |
| P10153 | -1,62 | Negative | 3,36 |
| O75144 | -1,63 | Negative | 3,72 |
| Q01973 | -1,63 | Negative | 5,42 |
| P35908 | -1,63 | Negative | 3,20 |
| Q9UNN8 | -1,64 | Negative | 4,34 |
| Q07507 | -1,64 | Negative | 4,28 |
| P04899 | -1,65 | Negative | 5,41 |
| P55287 | -1,65 | Negative | 4,26 |
| P56537 | -1,66 | Negative | 5,07 |
| P19440 | -1,66 | Negative | 3,75 |
| P63000 | -1,66 | Negative | 4,48 |
| Q9UKU9 | -1,67 | Negative | 5,25 |
| P26992 | -1,67 | Negative | 6,09 |
| P24821 | -1,68 | Negative | 5,45 |
| O00592 | -1,68 | Negative | 3,19 |
| P10253 | -1,68 | Negative | 6,72 |
| Q7Z3B1 | -1,68 | Negative | 4,13 |
| P07858 | -1,70 | Negative | 4,99 |
| P07911 | -1,71 | Negative | 5,28 |
| P08294 | -1,71 | Negative | 4,92 |
| P19835 | -1,72 | Negative | 4,07 |
| Q9HD42 | -1,72 | Negative | 2,98 |
| P08571 | -1,72 | Negative | 3,03 |
| Q15375 | -1,72 | Negative | 7,75 |
| Q9Y2E5 | -1,72 | Negative | 5,31 |
| Q92859 | -1,73 | Negative | 6,10 |
| Q12805 | -1,73 | Negative | 4,70 |
| P07998 | -1,74 | Negative | 3,02 |
| Q15907 | -1,74 | Negative | 7,36 |
| P01591 | -1,74 | Negative | 3,88 |
| Q15746 | -1,74 | Negative | 3,74 |
| Q9HBB8 | -1,75 | Negative | 3,18 |
| P05091 | -1,76 | Negative | 4,74 |
| O43451 | -1,76 | Negative | 3,86 |
| P05452 | -1,76 | Negative | 4,84 |
| P63092; Q5JWF2 | -1,76 | Negative | 8,03 |
| Q9UBX5 | -1,76 | Negative | 5,63 |
| Q99574 | -1,77 | Negative | 6,23 |
| O15197 | -1,77 | Negative | 7,21 |
| P98164 | -1,78 | Negative | 5,99 |
| A0A075B6I9 | -1,79 | Negative | 3,21 |
| P08519 | -1,79 | Negative | 2,74 |
| P14543 | -1,80 | Negative | 8,06 |
| P12429 | -1,81 | Negative | 3,26 |
| Q8IUL8 | -1,82 | Negative | 5,10 |
| Q08174 | -1,83 | Negative | 5,27 |
| Q9UIB8 | -1,83 | Negative | 5,14 |
| P43121 | -1,84 | Negative | 7,49 |
| Q8N6C8 | -1,86 | Negative | 6,47 |
| Q8WZ75 | -1,86 | Negative | 5,74 |
| Q9UBD6 | -1,86 | Negative | 3,71 |
| P58499 | -1,87 | Negative | 7,58 |
| Q9BXP8 | -1,87 | Negative | 6,15 |
| Q9UBI6 | -1,88 | Negative | 2,42 |
| Q9Y6R7 | -1,88 | Negative | 2,47 |
| Q8IV08 | -1,89 | Negative | 7,72 |
| P40189 | -1,89 | Negative | 8,51 |
| Q5T011 | -1,89 | Negative | 2,96 |
| P61916 | -1,90 | Negative | 5,15 |
| O75339 | -1,90 | Negative | 5,41 |
| Q03403 | -1,91 | Negative | 3,91 |
| P06703 | -1,94 | Negative | 3,30 |
| O43278 | -1,94 | Negative | 10,56 |
| Q9UJX4 | -1,95 | Negative | 2,71 |
| P05451 | -1,95 | Negative | 2,74 |
| Q7Z3E2 | -1,96 | Negative | 4,84 |
| P54760 | -1,97 | Negative | 9,31 |
| P98172 | -1,98 | Negative | 6,59 |
| Q8IYS5 | -1,99 | Negative | 4,65 |
| P50895 | -1,99 | Negative | 6,86 |
| P12830 | -2,01 | Negative | 5,57 |
| P11279 | -2,02 | Negative | 5,66 |
| Q14956 | -2,02 | Negative | 4,90 |
| Q15828 | -2,04 | Negative | 2,66 |
| P01833 | -2,04 | Negative | 10,27 |
| P80188 | -2,05 | Negative | 4,43 |
| P02751 | -2,05 | Negative | 6,26 |
| Q96RW7 | -2,09 | Negative | 6,41 |
| Q6UVK1 | -2,11 | Negative | 7,77 |
| Q9GZX9 | -2,13 | Negative | 3,20 |
| P14384 | -2,14 | Negative | 8,54 |
| O00533 | -2,15 | Negative | 7,84 |
| Q969P0 | -2,16 | Negative | 6,44 |
| Q7KYR7; Q8WVV5 | -2,17 | Negative | 5,58 |
| P19961 | -2,18 | Negative | 4,81 |
| Q9P121 | -2,22 | Negative | 8,78 |
| Q6UXG3 | -2,22 | Negative | 3,94 |
| P08174 | -2,22 | Negative | 5,44 |
| Q9BX67 | -2,22 | Negative | 5,98 |
| O60494 | -2,23 | Negative | 8,65 |
| Q9UMR5 | -2,26 | Negative | 8,14 |
| P04745; P0DTE7; P0DTE8; P0DUB6 | -2,27 | Negative | 7,08 |
| Q96NY8 | -2,30 | Negative | 7,71 |
| O95716 | -2,32 | Negative | 6,10 |
| P08637 | -2,33 | Negative | 5,30 |
| P06870 | -2,33 | Negative | 5,05 |
| Q8NFZ8 | -2,34 | Negative | 7,58 |
| P0DP57 | -2,35 | Negative | 5,12 |
| P54753 | -2,37 | Negative | 12,14 |
| Q8N6Q3 | -2,37 | Negative | 6,42 |
| Q9UN70 | -2,39 | Negative | 8,09 |
| Q14982 | -2,42 | Negative | 8,03 |
| P15328 | -2,44 | Negative | 6,89 |
| P62879 | -2,46 | Negative | 8,12 |
| P78324 | -2,47 | Negative | 8,97 |
| P13987 | -2,51 | Negative | 6,63 |
| P30530 | -2,52 | Negative | 7,16 |
| Q9BQ51 | -2,54 | Negative | 7,30 |
| O75594 | -2,54 | Negative | 8,31 |
| Q8NBJ4 | -2,85 | Negative | 7,86 |
| Q9NYU1; Q9NYU2 | -3,09 | Negative | 9,70 |
| Q9H299 | -3,10 | Negative | 4,12 |

**Supplementary Table 2. Top 20 proteins identified from the classifier**

| **Common^a^** | **Unique^b^** |
| --- | --- |
| P01009 (SERPINA1) | Q9UMR5 (PPT2) |
| P43652 (AFM) | Q9Y5Y6 (ST14) |
| P02768 (ALB) | Q12841 (FSTL1) |
| P78386 (KRT85) | O15197 (EPHB6) |
| P00450 (CP) |  |
| Q9Y625 (GPC6) |  |
| P01008 (SERPINC1) |  |
| P02787 (TF) |  |
| P02748 (C9) |  |
| P02766 (TTR) |  |
| P01833 (PIGR) |  |
| O75882 (ATRN) |  |
| P02774 (GC) |  |
| Q9NYU1 (UGGT2) |  |
| Q92859 (NEO1) |  |
| P14543 (NID1) |  |

Gene names are shown in parentheses.
^a^Overlap with the list differentially abundant proteins.
^b^Unique to the machine learning algorithm.
